# Unexpected plasticity in the life cycle of *Trypanosoma brucei*

**DOI:** 10.1101/717975

**Authors:** Sarah Schuster, Ines Subota, Jaime Lisack, Henriette Zimmermann, Christian Reuter, Brooke Morriswood, Markus Engstler

**Affiliations:** Department of Cell and Developmental Biology, Biocentre, University of Wuerzburg; University of Wuerzburg, Germany

## Abstract

African trypanosomes cause sleeping sickness in humans and nagana in cattle. These unicellular parasites are transmitted by the bloodsucking tsetse fly. In the mammalian host’s circulation, tissues, and interstitium, at least two main life cycle stages exist: slender and stumpy bloodstream stages. Proliferating slender stage cells differentiate into cell cycle-arrested stumpy stage cells at high population densities. This developmental stage transition occurs in response to the quorum sensing factor SIF (stumpy induction factor), and is thought to fulfil two main functions. First, it auto-regulates the parasite load in the host. Second, the stumpy stage is regarded as pre-adapted for tsetse fly infection and the only stage capable of successful vector transmission. Here, we show that proliferating slender stage trypanosomes are able to complete the complex life cycle in the fly as successfully as the stumpy stage, and that a single parasite is sufficient for productive infection. Our findings not only propose a revision to the traditional rigid view of the trypanosome life cycle, but also suggest a solution to a long-acknowledged paradox in the transmission event: parasitaemia in chronic infections is characteristically low, and so the probability of a tsetse ingesting a stumpy cell during a bloodmeal is also low. The finding that proliferating slender parasites are infective to tsetse flies helps shed light on this enigma.

## Introduction

Trypanosomes are among the most successful parasites. These flagellated protists infect all vertebrate classes, from fish to mammals, and can cause devastating diseases. African trypanosomes, which are transmitted by the tsetse fly, are the agents of nagana in livestock and sleeping sickness in humans (Bruce, London School of, & Tropical, 1895). The most intensively-studied African trypanosome species is *Trypanosoma brucei*, which in the past decades has emerged as a genetic and cell biological model parasite. The general life cycle of *T. brucei* was elucidated more than a century ago. As part of this life cycle, the trypanosomes undergo a full developmental program in the tsetse fly in order to become infective (Koch, 1909). This finding, made by Kleine in 1909, showed that transmission was not a purely mechanical event (Kleine, 1909). Kleine subsequently found that the life cycle in the fly could take up to several weeks to complete, a discovery that was shortly afterwards confirmed by Bruce (Bruce, Hamerton, Bateman, & Mackie, 1909). More details of the general life cycle of *Trypanosoma brucei* were then elucidated by Robertson in 1913, with several key observations concerning the transmission event (Robertson & Bradford, 1913). Subsequent work has resulted in a detailed picture of the passage through the fly, beginning with the ingestion of trypanosomes in an infected bloodmeal (Rotureau & Van Den Abbeele, 2013). After entering through the tsetse proboscis, the infected blood is either held for a short time in the crop, which acts as a storage site, allowing tsetse to drink more blood per meal, or is passed directly to the midgut. Upon entering the tsetse midgut, the trypanosomes differentiate into the proliferative procyclic stage. Once established in the midgut, the parasites must pass the peritrophic matrix, a protective sleeve that separates the bloodmeal from midgut tissue. To do this, the parasites swim up the endotrophic space to the proventriculus, the site of peritrophic matrix synthesis, where they are able to cross to the ectotrophic space. After having crossed the peritrophic matrix and entered the ectroperitrophic space, procyclic trypanosomes may either further colonize the ectotrophic anterior midgut, becoming the cell-cycle arrested mesocyclic stage, or continue directly to the proventriculus. In the proventriculus, trypanosomes further develop into the long, proliferative epimastigote stage (Rose et al., 2020). The epimastigotes then swim from the proventriculus to the salivary glands, while undergoing an asymmetric division to generate a long and a short daughter cell. Once in the salivary gland, the long daughter cell dies while the small one attaches via its flagellum to the salivary gland epithelium (Vickerman, 1969). The attached epimastigotes are proliferative, producing either more attached epimastigote daughter cells or freely swimming, cell cycle-arrested metacyclic trypanosomes. As early as 1911, it was clear that the metacyclic stage (at that time called metatrypanosomes) is the only mammalian-infective stage (Bruce, Hamerton, Bateman, & Mackie, 1911).

In the mammalian host, trypanosomes have been found in many different organs, including brain tissue, skin, and fat, but are hard to study experimentally (Capewell et al., 2016; Goodwin, 1970; Krüger, Schuster, & Engstler, 2018; Trindade et al., 2016). The two main stages found in the bloodstream, and the best-characterised experimentally, are the proliferating slender bloodstream stage and the cell cycle-arrested stumpy bloodstream stage (Krüger et al., 2018; Keith R. Matthews, Ellis, & Paterou, 2004; Vickerman, 1985). The stumpy stage is formed in response to quorum sensing of the stumpy induction factor (SIF), a signal produced by slender bloodstream trypanosomes (Vassella, Reuner, Yutzy, & Boshart, 1997). As the stumpy stage only survives for 2-3 days after formation, the generation of stumpy parasites is thought to control the burden the parasites impose on the host (Turner, Aslam, & Dye, 1995). The SIF pathway that controls the slender-to-stumpy transition has been detailed down to the molecular level, with the *protein associated with differentiation* (PAD1) as the first recognised molecular marker for the stumpy stage (Dean, Marchetti, Kirk, & Matthews, 2009; Mony & Matthews, 2015). More recently, it was also shown that the stumpy pathway can be triggered independently of SIF, though the extent to which this occurs in the general population remains unclear (Batram, Jones, Janzen, Markert, & Engstler, 2014; Zimmermann et al., 2017). Besides its proposed role in controlling parasitaemia in the mammalian host, the stumpy stage has a second essential function in the trypanosome life cycle: it is believed to be the only life cycle stage that can infect the tsetse fly (Rico et al., 2013). Thus, arrest of the cell cycle and differentiation to the stumpy stage are presumed essential for developmental progression to the procyclic insect stage. As early as 1912, Robertson suggested that the short, stumpy bloodstream trypanosomes represent the fly-infective stage (Robertson, 1912). While this assumption was questioned several times throughout the 20^th^ century, the discovery of quorum sensing and SIF in the 1990s made it become generally accepted (Vassella et al., 1997). However, if stumpy trypanosomes are the only stage that can infect the fly, another problem arises. Chronic trypanosome infections are characterised by low blood parasitemia, meaning that the chance of a tsetse fly ingesting any trypanosomes, let alone short-lived stumpy ones is also very low (Frezil, 1971; Wombou Toukam, Solano, Bengaly, Jamonneau, & Bucheton, 2011). Mathematical models have been developed that aim to explain how the limited number of short-lived stumpy cells in the host blood and interstitial fluids can guarantee the infection of the tsetse fly, which is essential for the survival of the species (Capewell et al., 2019; MacGregor & Matthews, 2008; Seed & Black, 1999). The present study provides surprising new solutions to this problem. First, systematic quantification of infection efficiencies showed that very few trypanosomes are necessary to infect a tsetse fly, and in fact just one is sufficient. Second, and wholly unexpectedly, slender stages proved at least as competent at infecting flies as stumpy stages. These findings suggest greater flexibility in the life cycle than supposed, prompting a revision to the current rigid view of the process.

## Results

### A single trypanosome is sufficient for infection of a tsetse fly

Slender and stumpy bloodstream stage trypanosomes can be distinguished based on cell cycle, morphological, and metabolic criteria. The genome of the single mitochondrion (kinetoplast, K) and the cell nucleus (N) can be readily visualized using DNA stains, and their prescribed sequence of replication (1K1N, 2K1N, 2K2N) allows cell cycle stage to be inferred (Sherwin & Gull, 1989). Slender cells are found in all three K/N ratios, while stumpy cells, which are cell cycle-arrested, are found only as 1K1N cells (Fig. 1A). Expression of the *protein associated with differentiation 1* (PAD1) is accepted as a marker for development to the stumpy stage (Dean et al., 2009). Cells expressing an NLS-GFP reporter fused to the 3’ UTR of the PAD1 gene (GFP:PAD1^UTR^) will have GFP-positive nuclei when the PAD1 gene is active. Hence, slender cells are GFP-negative; stumpy cells are GFP-positive (Fig. 1A). The validity of the NLS-GFP reporter as an indicator for the activation of the PAD1 pathway (Batram et al., 2014; Zimmermann et al., 2017) was corroborated by co-staining with an antibody against the PAD1 protein (Supplementary Fig. 1). We have previously shown that stumpy cells can be formed independently of high cell population density ectopic expression of a second variant surface glycoprotein (VSG) isoform, a process that mimics one of the pathways involved in trypanosome antigenic variation (Batram et al., 2014; Cross, 1975; Hertz-Fowler et al., 2008; Zimmermann et al., 2017). These so-called expression site (ES)-attenuated stumpy cells can complete the developmental cycle in the tsetse fly (Zimmermann et al., 2017). It remained an open question whether this occurred with the same efficiency as with

**Figure 1.**
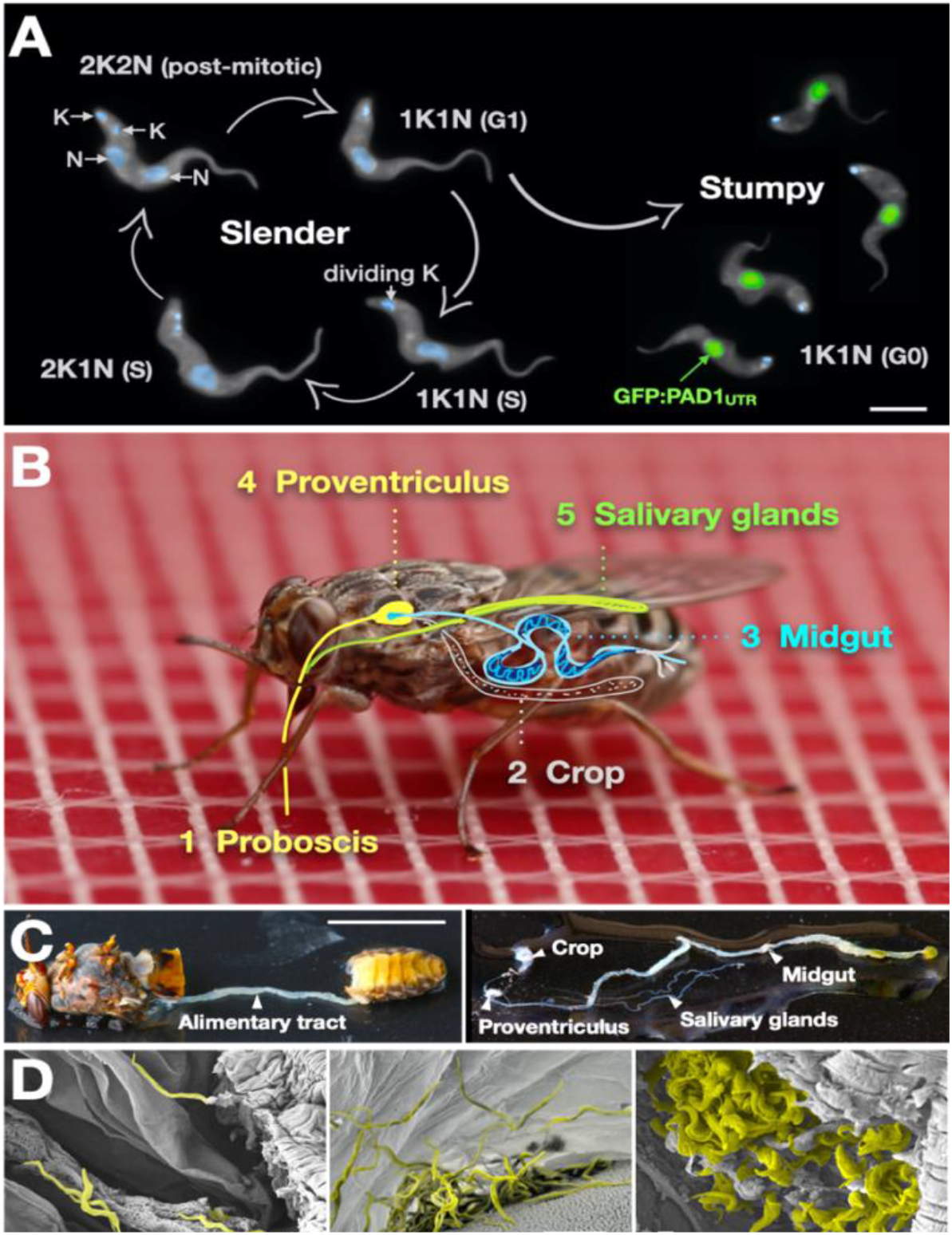
Slender trypanosomes can complete the entire life cycle in the tsetse fly vector. (A) Cell cycle (G1/S/post-mitotic), morphology, and differentiation of bloodstream form (mammalian-infective stage) trypanosomes. Proliferation of slender trypanosomes is detectable by duplication and segregation of the mitochondrial genome (kinetoplast, K) and nuclear DNA (N) over time. Quorum sensing causes cell cycle arrest (G0) and expression of the stumpy marker PAD1. Images are false-coloured, maximum intensity projections of deconvolved 3D stacks. The green colour indicates the nuclear GFP:PAD1^UTR^ fluorescence, the DAPI-stained kinetoplast and nucleus are shown in light blue, and the AMCA-sulfo-NHS-labelled parasite cell surface is shown in gray. Scale bar: 5 μm. (B) Trypanosome infections of tsetse flies were achieved via bloodmeal, which consists typically of 20 μl, through a silicone membrane. To complete infection in a tsetse fly after an infective bloodmeal, trypanosomes first travel to the midgut, followed by the proventriculus, and finally must reach the salivary glands. The corresponding video is available in the Supplementary information (Supplementary Video 1). (C) The first panel depicts a dissected, infected tsetse fly for explantation of the alimentary tract. The second panel shows the explanted alimentary tract of the tsetse, with the different subcompartments labelled. Scale bar: 5 mm.(D) Scanning electron micrograph of a typical trypanosome infection of the tsetse midgut, proventriculus, and salivary glands. Parasites are false-coloured yellow. Scale bar: 1 μm.

SIF-produced stumpy cells. Therefore, we quantitatively compared the transmission competence of stumpy populations generated by either SIF treatment or through ES-attenuation. Tsetse flies *(Glossina morsitans morsitans’)* were infected via membrane feeding (Fig. 1B; Supplementary Video 1) with defined numbers of pleomorphic stumpy trypanosomes, capable of completing the entire developmental cycle. This cycle includes entrance through the proboscis, passage through the crop, establishing infections in the midgut, proventriculus, and finally the salivary glands (Fig. 1B). Two transgenic trypanosome cell lines, both of which contained the GFP:PAD1^UTR^ reporter construct, were used. One was subjected to tetracycline-induced, ectopic VSG expression to drive ES attenuation (Table 1, lines i-iii, Stumpy^ES^) (Zimmermann et al., 2017). The other was treated with stumpy induction factor (Table 1, lines iv-vi, Stumpy^SIF^). Both treatments resulted in expression of the GFP:PAD1^UTR^ reporter and rapid differentiation to the stumpy stage. The resulting stumpy populations were fed to tsetse flies at concentrations ranging from 120,000 to 10 cells/ml. A feeding tsetse typically ingests about 20 μl of blood (Gibson & Bailey, 2003), meaning that, on average, between 2,400 and 0.2 trypanosomes were ingested per bloodmeal (Table 1, rows i-vi, column 2, Total). The trypanosomes had previously been scored for expression of the GFP:PAD1^UTR^ reporter to confirm their identity as the stumpy stage (Table 1, columns 3-4). To analyse the infections, we carried out microscopic analyses of explanted tsetse digestive tracts (Fig. 1C). The dissection of the flies was done 5-6 weeks post infection. The presence of mammal-infective, metacyclic trypanosomes in explanted tsetse salivary glands indicated the completion of the life cycle inside the tsetse. Remarkably, the uptake, on average, of two stumpy parasites of either cell line produced robust infections of tsetse midgut (MG), proventriculus (PV), and salivary glands (SG) (Fig. 1D; Table 1, columns 5-7). Ingestion, on average, of even a single stumpy cell was sufficient to produce salivary gland infections in almost 5% of all tsetse (Table 1, row v). When the stumpy parasite number was further reduced to 0.2 cells on average per bloodmeal, meaning every 5th fly would receive a stumpy cell, 0.9% of flies still acquired salivary gland infections (Table 1, row vi). As a measure of the incidence of life cycle completion in the tsetse fly, we calculated the transmission index (TI) for each condition. The TI has been defined as the ratio of salivary gland to midgut infections and hence, it is a measure for successful passage through the second part of the trypanosome tsetse cycle (Fig. 1B, 4-5)(Peacock, Ferris, Bailey, & Gibson, 2012). We found that for flies infected with 2 trypanosomes on average, the TI was comparable between SIF-induced (TI = 0.29) and ES-induced (TI = 0.31) stumpy trypanosomes (Table 1, rows iii-iv; Fig. 2). A similar TI of 0.23 was observed in flies ingesting on average 1 trypanosome (Table 1, row v; Fig. 2). Thus, our data not only clearly show that SIF- and ES-induced stumpy parasites are equally efficient in completing the weeks-long, multi-step fly cycle, but also that a single stumpy cell is sufficient to produce a mature fly infection. While this may seem comparable with an observation that has been made before for *Trypanosoma congolense*(Maudlin & Welburn, 1989), the migration through the fly differs between the two species: *T. brucei* infects the salivary glands, while *T. congolense* infects the proboscis. The tsetse fly, however, is much more susceptible to infections with *T. congolense* than with *T. brucei*, with a nearly 5-fold increase in percent *T. congolense* proboscis infections as compared to *T. brucei* salivary gland infections. As the authors used GSH and NAG to boost T. brucei infections, the 5-fold difference is actually a lower estimate. (Peacock et al., 2012). Our results demonstrate that very low numbers of *T. brucei* stumpy cells can also successfully establish mature tsetse fly infections.

**Table 1.** Slender trypanosomes can complete the entire tsetse infection cycle, and a single parasite is sufficient for tsetse passage. The flies were infected with either stumpy or slender trypanosomes. Stumpy trypanosomes were generated by induction of expression site attenuation (ES), or SIF-treatment (SIF). MG, midgut infection; PV, proventriculus infection; SG, salivary gland infection; TI, transmission index (SG/MG); n, number of independent fly infection experiments.

| Number of trypanosomes per blood meal |  |  |  |  | Tsetse fly infection (%) |  |  |  |  |  |  |  |
| --- | --- | --- | --- | --- | --- | --- | --- | --- | --- | --- | --- | --- |
|  | 1 | 2 | 3 | 4 | 5 | 6 | 7 | 8 | 9 | 10 | 11 | n |
|  |  | Total | PAD1-negative | PAD1-positive | MG | PV | SG | TI | Tsetse infected | Tsetse dissected | Sex ratio (♀/♂) |  |
| i | Stumpy<sup>ES</sup> | 2400 | 105 | 2295 | 19.3 | 18.1 | 12 | 0.63 | 107 | 83 | 1.04 | 3 |
| ii |  | 20 | 1 | 19 | 25.3 | 22.9 | 9.6 | 0.38 | 115 | 83 | 0.39 | 4 |
| iii |  | 2 | 0 | 2 | 14.6 | 14.6 | 4.5 | 0.31 | 110 | 89 | 0.96 | 4 |
| iv | Stumpy<sup>SIF</sup> | 2 | 0 | 2 | 38.8 | 29.3 | 11.2 | 0.29 | 120 | 116 | 1.70 | 2 |
| v |  | 1 | 0 | 1 | 20.2 | 18.3 | 4.6 | 0.23 | 122 | 109 | 0.60 | 2 |
| vi |  | 0.2 | 0 | 0.2 | 7.5 | 4.7 | 0.9 | 0.13 | 114 | 107 | 1.28 | 2 |
| vii | Slender<sup>ES</sup> | 20 | 19.16 | 0.84 | 22.5 | 22.5 | 5.0 | 0.22 | 104 | 80 | 0.67 | 3 |
| viii |  | 2 | 1.93 | 0.07 | 13.2 | 13.2 | 7.9 | 0.60 | 100 | 76 | 0.96 | 3 |
| ix | Slender<sup>SIF</sup> | 2 | 1.93 | 0.07 | 7.8 | 7.8 | 4.7 | 0.60 | 156 | 129 | 1.44 | 3 |
| x |  | 1 | 0.99 | 0.01 | 4.7 | 4.1 | 2.0 | 0.44 | 383 | 343 | 1.01 | 6 |
| xi |  | 0.2 | 0.002 | 0.198 | 0.9 | 0.9 | 0.0 | 0.00 | 130 | 114 | 1.35 | 2 |
| xii | Slender<sup>naive</sup> | 2 | 1.971 | 0.029 | 6.3 | 3.6 | 2.7 | 0.43 | 350 | 331 | 0.96 | 6 |
| xiii | Monomorph | 2400 | 2395 | 5 | 11.0 | 0.6 | 0.0 | 0.00 | 186 | 155 | 0.64 | 3 |
| xiv |  | 20 | 20 | 0 | 1.4 | 0.0 | 0.0 | 0.00 | 216 | 127 | 0.76 | 3 |
| xv |  | 2 | 2 | 0 | 2.5 | 0.0 | 0.0 | 0.00 | 298 | 257 | 0.87 | 3 |

### Proliferating slender bloodstream stage trypanosomes infect the insect vector with comparable efficiency as cell cycle-arrested stumpy bloodstream stage parasites

Originally intended as a control experiment with an easily predictable (negative) outcome, we infected tsetse flies with proliferating PAD1-negative slender trypanosomes from the two pleomorphic cell lines used (Table 1, rows vii-xi). Unexpectedly, we found that slender parasites were not only viable in the midgut, but also infected the proventriculus and the salivary glands. (Table 1, rows vii-xi). Even one slender parasite was sufficient to establish solid midgut infections, proving that slender and stumpy parasites are, in principle, equally viable in the tsetse midgut. The infection efficiency when the flies were fed with either 20 stumpy trypanosomes or 20 pleomorphic slender trypanosomes was similar (Table 1, compare TI in column 8 for rows ii and vii). When flies were fed with an average of 2 slender parasites each, the TI was actually higher for slender cells (0.60) than for stumpy cells (0.31) (Fig. 2). This TI of 0.60 was identical for both populations of slender cells (Fig. 2). Next, when given, on average, just one PAD1-negative slender cell per bloodmeal, parasite infections were still established in the midgut, proventriculus, and salivary glands with incidences of 4.7%, 4.1%, and 2.0% respectively, at a TI of 0.44 (Table 1, row x; Fig. 2). In order to be absolutely sure that slender trypanosomes can passage through the tsetse, we repeated the experiment with naïve slender parasites that had been freshly differentiated from insect-derived metacyclic trypanosomes, i.e. cells that had just restarted the mammalian life cycle stage (Table 1, row xii). Infections with, on average, two freshly-differentiated slender trypanosomes per bloodmeal revealed 6.3% midgut and 2.7% salivary gland infections. The transmission index was 0.43. This important control formally ruled out that cultivated slender cells had undergone any kind of gain-of-function adaptation in culture that made them transmission-competent.

**Figure 2.**
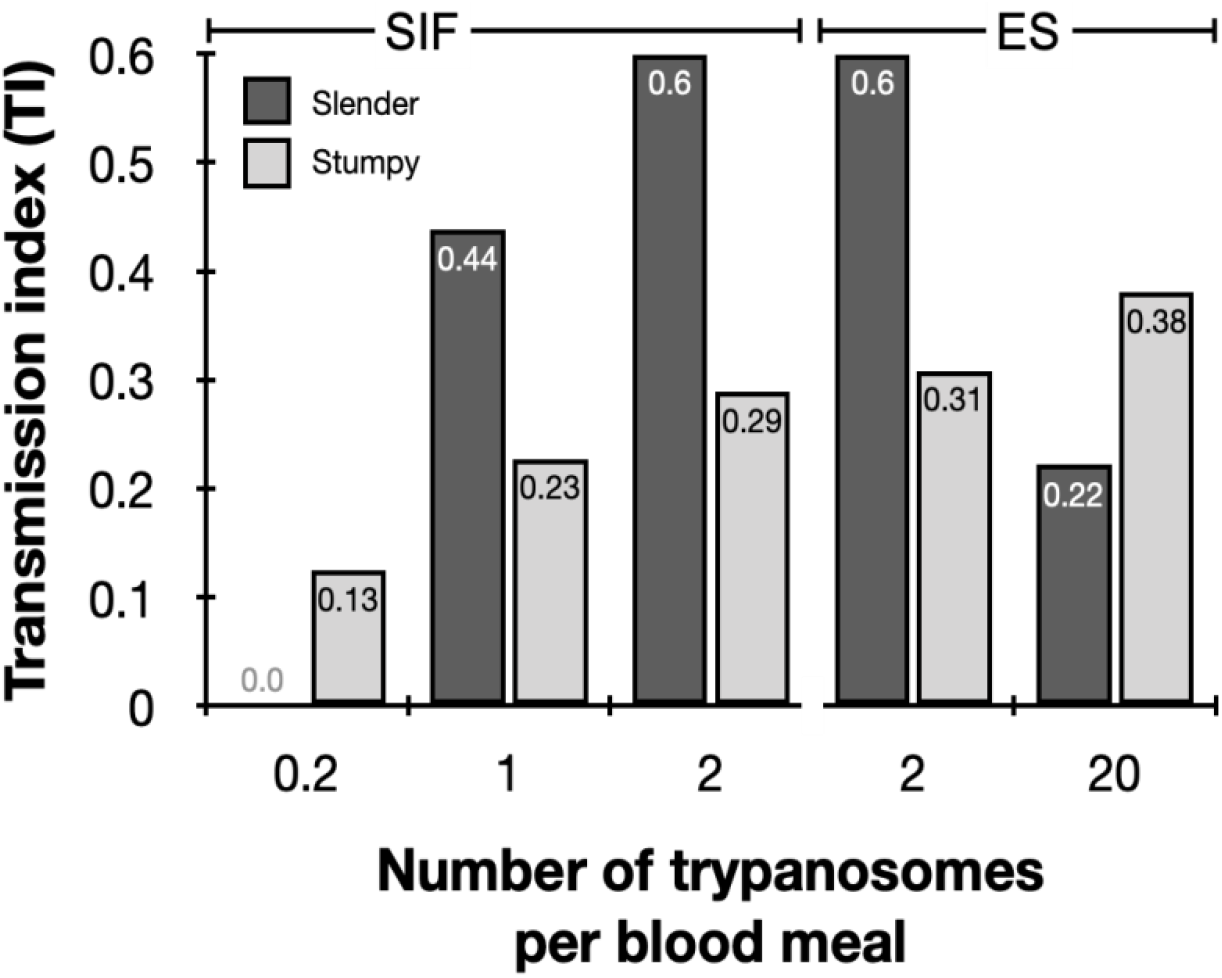
Graphical representation of the transmission index TI (SG/MG) of slender (dark gray) and stumpy (light gray) trypanosomes at different numbers per bloodmeal (data reproduced from Table 1, column 8). A high TI indicates successful completion of the life cycle in the tsetse vector. At low infective doses, slender trypanosomes had a higher TI compared to stumpy parasites. There was no difference between stumpy parasites generated by SIF-treatment (SIF) or expression site attenuation (ES).

As another control for the slender infection experiments, tsetse infections were carried out using a monomorphic slender trypanosome strain, i.e. one that had lost the capacity of differentiating to the stumpy stage (Table 1, rows xiii - xv). Monomorphic trypanosomes are known to be – in principle - able to infect the tsetse midgut, but they are incapable of completing the developmental cycle in the fly (Herder et al., 2007; Peacock, Ferris, Bailey, & Gibson, 2008). As expected, no salivary gland infections were seen using these cells, even at high infection numbers. Interestingly, we found that even two monomorphic slender parasites can establish a fly midgut infection (Table 1, row xv). Thus, infection of the tsetse midgut is independent of the capacity for developmental progression and the infective dose, and it does not require the stumpy life cycle stage. This finding also challenges the assumption that slender parasites are selectively eliminated from the parasite population and that only stumpy trypanosomes can survive the harsh conditions thought to prevail within the tsetse crop and midgut (Nolan, Rolin, Rodriguez, Van Den Abbeele, & Pays, 2000).

The ES-attenuated cells showed similar midgut, proventriculus, and salivary gland infection incidence as either the stumpy or slender stage (Table 1, rows ii-iii and vii-viii). The SIF-induced stumpy cells, however, appeared more effective in establishing midgut infections than their slender counterparts (Table 1, rows iv-vi and ix-xi). This result could be interpreted as stumpy trypanosomes being more successful in the tsetse fly, but this is a conclusion that is clearly not supported by our data. First, the infections with 1-2 slender cells produced higher TI values than those with the same numbers of stumpy cells (Fig. 2). This suggests that the proliferative slender cells are actually more capable of progressing from a midgut infection to a salivary gland one, and thus have at least comparable overall developmental competence to the stumpy stage. Second, the lack of correlation between infective dose and midgut infections underlines the importance of the TI as a relative measure. What is biologically relevant is not the initiation of infection but the completion of the tsetse passage. In summary, our experiments not only establish that a single *T. brucei* (either slender or stumpy) parasite can infect the tsetse fly, but also proves that slender cells can efficiently complete the passage through the tsetse fly.

### In the tsetse midgut, dividing slender bloodstream stage parasites activate the PAD1 pathway and differentiate to the procyclic insect stage without arresting the cell cycle

To determine how pleomorphic slender trypanosomes manage to establish infections, we observed the early events following trypanosome ingestion by tsetse flies (Supplementary Video 2). The canonical version of events is that ingested stumpy (i.e. PAD1-positive) cells reactivate the cell cycle, begin to express the EP procyclin protein on their cell surface, and differentiate to the procyclic life cycle stage(Dean et al., 2009; K R Matthews & Gull, 1994; Mowatt & Clayton, 1987; Richardson, Beecroft, Tolson, Liu, & Pearson, 1988; Roditi et al., 1989; Ziegelbauer & Overath, 1990). We infected tsetse with pleomorphic trypanosomes, which not only contained the stumpy-specific GFP:PAD1^UTR^ marker, but also encoded an EP1:YFP fusion (Fig. 3)(Engstler & Boshart, 2004). In this way, the onset of stumpy development was observable as GFP fluorescence in the nucleus, and further differentiation to the procyclic life cycle stage as YFP fluorescence on the parasite cell surface. In addition, the cell cycle status (K/N counts, see Fig. 1A), morphology, and the characteristic motile behavior of the trypanosomes were also assessed as criteria of developmental progress. In total, 114 tsetse flies (57 male and 57 female) were dissected after at least six independent infections with either 12,000 slender or stumpy parasites each. These high initial parasite numbers allowed the microscopic analysis of individual living slender (n = 1845) and stumpy trypanosomes (n = 1237) within the convoluted microenvironment of midgut explants (Schuster et al., 2017). As early as 2-4 h post-infection with slender trypanosomes, a few (0.8%) 2K1N dividing trypanosomes with a nuclear PAD1 signal could be observed (Fig. 4). After 8-10 hours however, half (38.3+6.8+5.3=50.4%) of all trypanosomes in the explants were PAD1-positive (Fig. 4, bar chart shows summed cell cycle category values for PAD1-positive cells). After 24 hours, 84.3% (56.3+15.0+13.0) of the parasites expressed PAD1. Of these, 9.8% had already initiated developmental progression to the procyclic insect stage, as evidenced by EP1:YFP fluorescence on their cell surface (Fig. 5). At 48-50 h post-infection with slender trypanosomes, virtually the entire trypanosome population (91.8%) expressed PAD1, and almost one fifth (19.1%) of cells were EP1-positive (Fig. 5). To examine cell cycle progression, we counted the number of 1K1N, 2K1N, and 2K2N cells in the PAD1-positive and PAD1-negative slender cell populations (Fig. 4, Slender rows). Remarkably, 15-17 h post-infection, the majority of all replicating (i.e. 2K1N, 2K2N) cells were PAD1-positive (Fig. 4). No indication for a transient cell cycle arrest or intermittent impairment of cell cycle progression was observed. Over the duration of the experiment, PAD1-negative cells gradually decreased in numbers, while PAD1-positive slender cells at all cell cycle stages were increasingly observed (Fig. 4, dotted green and blue lines; Supplementary Video 2C). After two days, more than 90% of dividing trypanosomes were PAD1-positive. Thus, the PAD1 pathway was triggered in slender trypanosomes upon ingestion by the fly, and without prior cell cycle arrest.

**Figure 3.**
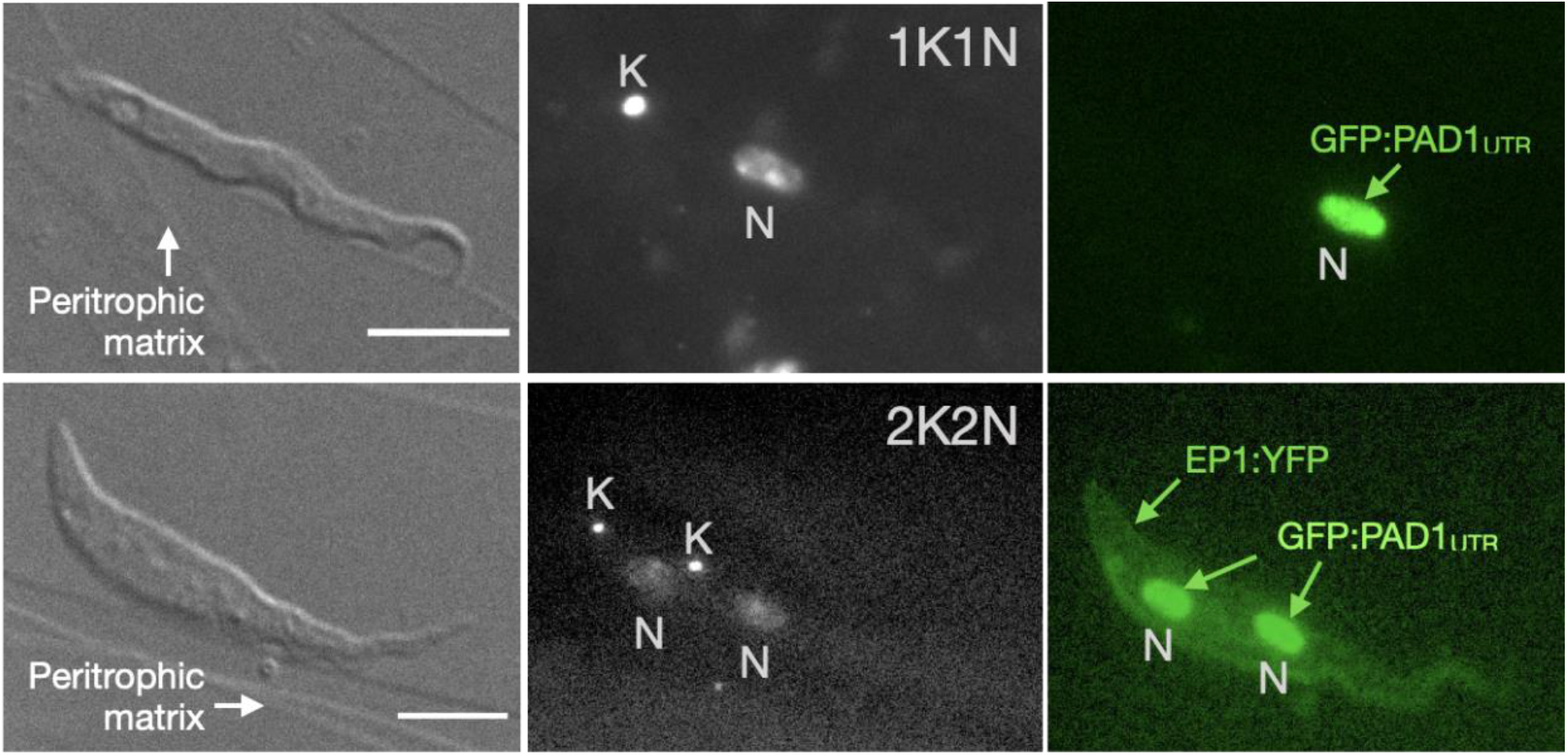
Exemplary images of procyclic trypanosomes in the tsetse explants 24 hours post infection with slender cells. Morphology (DIC panels, left), cell cycle status (DAPI label, middle panels) and expression of fluorescent reporters (right) were scored. Note that the upper panels show a cell with procyclic morphology that is nonetheless EP1:YFP negative, indicating that the EP1 signal underestimates the total numbers of procyclic cells in the population. Scale bar: 5 μm.

**Figure 4.**
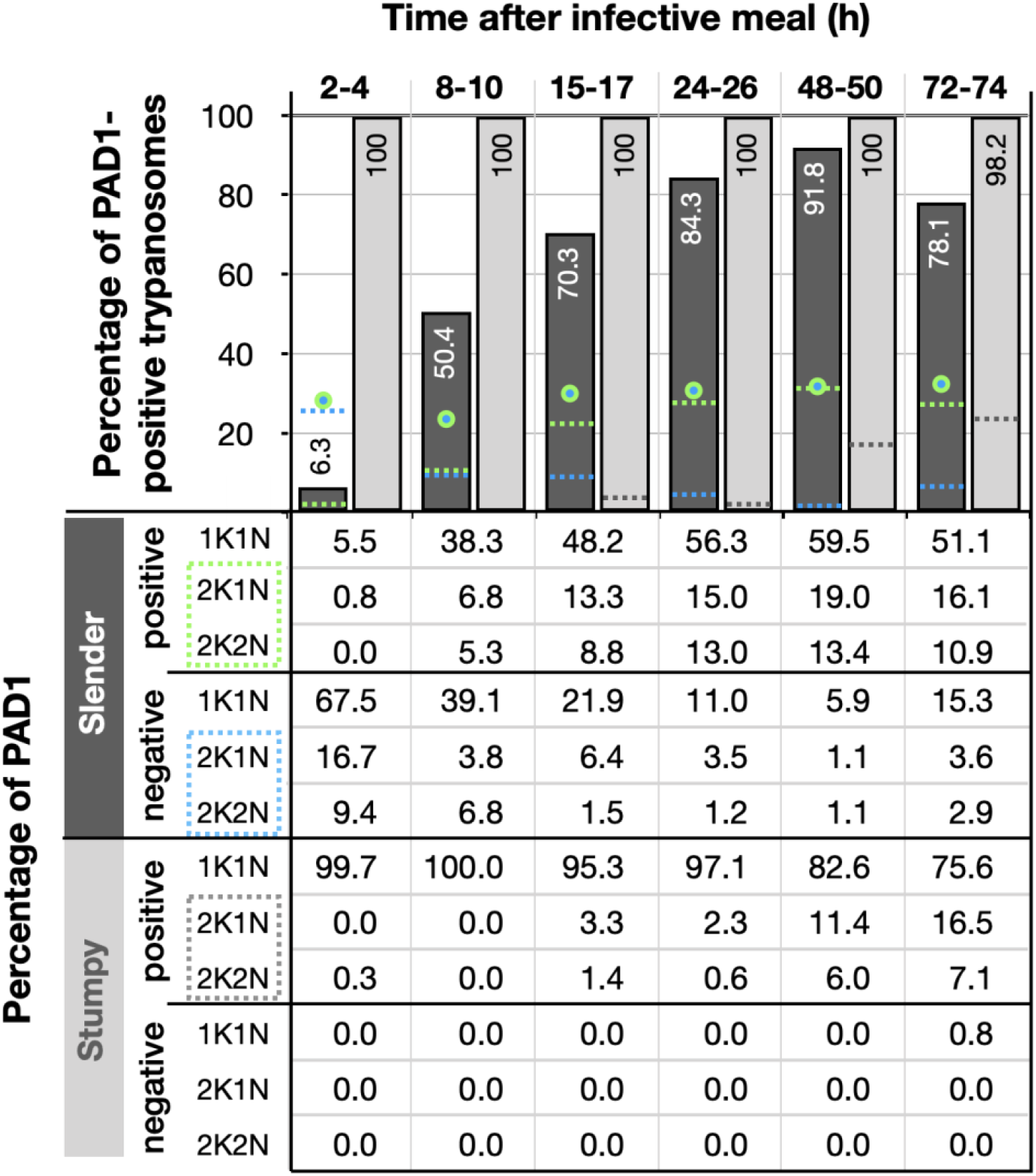
Slender trypanosomes activate the PAD1 pathway upon uptake by the tsetse fly, without cell cycle arrest. Tsetse flies were infected with either slender (3.6% PAD1-positive) or stumpy (100% PAD1-positive) trypanosomes. 72 (slender) or 42 (stumpy) flies were dissected (equal sex ratios) at different timepoints after infection. Experiments were done at least three times; data are presented as sample means. Living trypanosomes (>100 cells per time point) were microscopically analysed in the explants and scored for the expression of the fluorescent stumpy reporter GFP:PAD1^UTR^ in the nucleus. Stumpy cells (n=1237) are dark gray bars and slender cells (n=1845) are light gray bars. Slender and stumpy trypanosomes scored as PAD1-positive or -negative were also stained with DAPI, and the cell cycle position determined based on the configuration of kinetoplast (K) to nucleus (N) at the timepoints. Percentages of the population that were PAD1-positive and PAD-1 negative in the different cell cycle stages are indicated in the bottom table. The cell cycle stages are also displayed visually in top bar graph. As seen, while the total percentage of dividing slender cells remains constant over time (blue/green circles), the percentage of PAD-1 positive slender cells steadily increases (dotted green line) and the percentage of PAD-1 negative cells steadily decreases. This shows that slender cells can seamlessly turn on the PAD-1 pathway, without arresting in the cell cycle. Stumpy cells do not start having a normal cell cycle profile until 48 hours after tsetse uptake (dotted gray line - all PAD-1 positive), as the cells transition to the procyclic stage.

**Figure 5.**
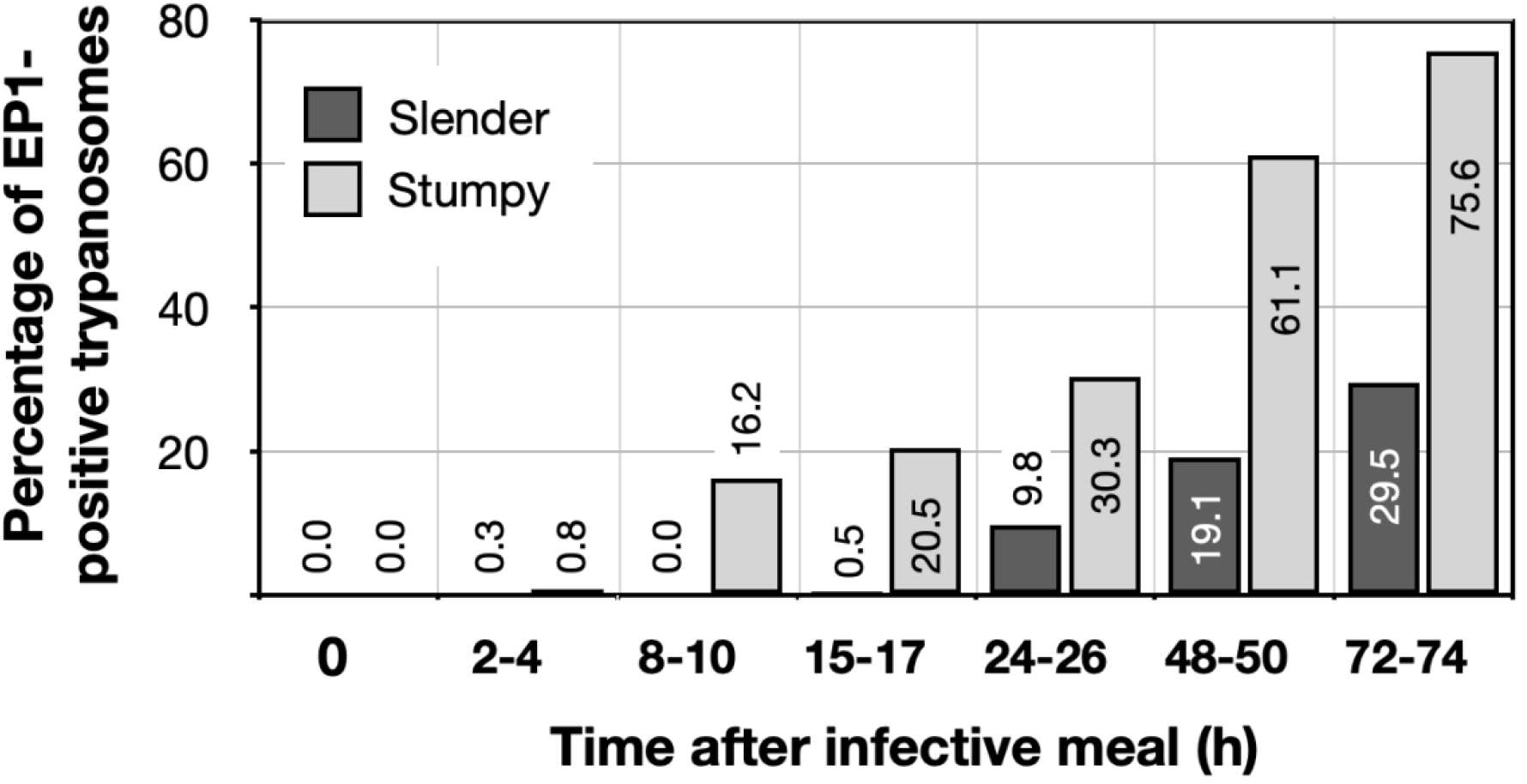
Slender trypanosomes differentiate to the procyclic life cycle stage in the tsetse fly without undergoing cell cycle arrest. Tsetse flies were infected with either slender (3.6% PAD1-positive) or stumpy (100% PAD1-positive) trypanosomes. 72 (slender) or 42 (stumpy) flies were dissected (equal sex ratios) at different timepoints after infection. Experiments were done at least three times; data are presented as sample means. Living trypanosomes (>100 cells per time point) were microscopically analysed in the explants and scored for the procyclic insect stage reporter EP1:YFP on the cell surface. Stumpy cells (n=1237) are shown by light gray bars and slender cells (n=1845) by dark gray bars.

In order to directly compare the kinetics of slender-to-procyclic development with that of stumpy stage trypanosomes, we fed flies with SIF-induced, PAD1-positive stumpy trypanosomes (Fig. 4, Stumpy rows). These cells remained as 1K1N cells in cell cycle arrest for the first day, and re-entered the cell cycle as procyclic parasites after 2 days. Four hours after uptake by the tsetse fly, stumpy trypanosomes started expressing EP1:YFP (Fig. 5). The fluorescent reporter was visible on 16.2 % of stumpy cells after 10 hours, showing that EP expression was initiated before release of cell cycle arrest. Uncoupling of EP surface expression from the commitment to differentiation has been reported before (Engstler & Boshart, 2004).

EP1:YFP expression in slender parasites lagged 12 hours behind stumpy cells, only becoming widespread after 24-26 hours (Fig. 5). Thus, the onset of EP1 expression was shifted, but the kinetics of differentiation were comparable in slender and stumpy parasites. Hence, activation of the PAD1 pathway also preceded developmental progression in slender cells. This means that expression of PAD1 is essential for differentiation to the insect stage, while cell cycle arrest is not. Of note, EP1 expression did not directly correlate with acquisition of procyclic morphology. At 24-26h, 9.8% of slender cells were EP1-positive (Fig. 5), but the EP1-negative cells frequently exhibited procyclic morphology (Fig. 3, upper panels). An example of a dividing (2K2N), PAD1-positive, EP1-positive cell is also shown (Fig. 3, lower panels; Supplementary Video 2D). Thus, it appears that a seamless developmental stage transition from the slender bloodstream stage to the procyclic insect stage took place, which was accompanied by the typical re-organization of the cytoskeleton and the concomitant switch of swimming styles (Heddergott et al., 2012; Schuster et al., 2017).

### Pleomorphic slender bloodstream stage trypanosomes can seamlessly differentiate to the procyclic insect stage without preceding cell cycle arrest in vitro

The factor(s) or condition(s) that trigger differentiation of bloodstream stage trypanosomes to the procyclic insect stage in the tsetse midgut are still ill-defined. In the laboratory, differentiation to the procyclic insect stage is routinely induced by the addition of *cis*-aconitate, a drop in glucose, and a temperature drop from 37°C to 27°C (Brun, Jenni, Schönenberger, & Schell, 1981; Czichos, Nonnengaesser, & Overath, 1986; Engstler & Boshart, 2004; Qiu et al., 2018; Ziegelbauer, Quinten, Schwarz, Pearson, & Overath, 1990) (Fig. 8).

We used this protocol to further investigate the developmental potential of cultivated pleomorphic slender bloodstream stage *in vitro* using the same cell lines and analysis as above (Fig. 6). Slender trypanosomes activated the PAD1 pathway rapidly after receiving the trigger, with 9.8% of all parasites being PAD1-positive within 2-4 hours, and 83.2% after 10 hours. PAD1 expression peaked after one day (98.3%), and declined thereafter (Fig. 6). Shortly after PAD1 reporter expression, EP1 appeared on the cell surface of 19.6% of all parasites within 8-10 hours, increasing to 98.3% after 3 days (Fig. 7). PAD1 and EP protein appearance on the cell surface was monitored throughout the timecourse using immunofluorescence (Supplemental Fig. 2). Throughout the timecourse, PAD1-positive 2K1N and 2K2N cells were continually observed, demonstrating that the PAD1-positive slender parasites did not arrest in the cell cycle, and continued dividing throughout *in vitro* differentiation to the procyclic stage (Fig. 6, Slender rows). After 3 days of *cis* aconitate treatment *in vitro*, slender trypanosomes had established a proliferating procyclic parasite population.

**Figure 6.**
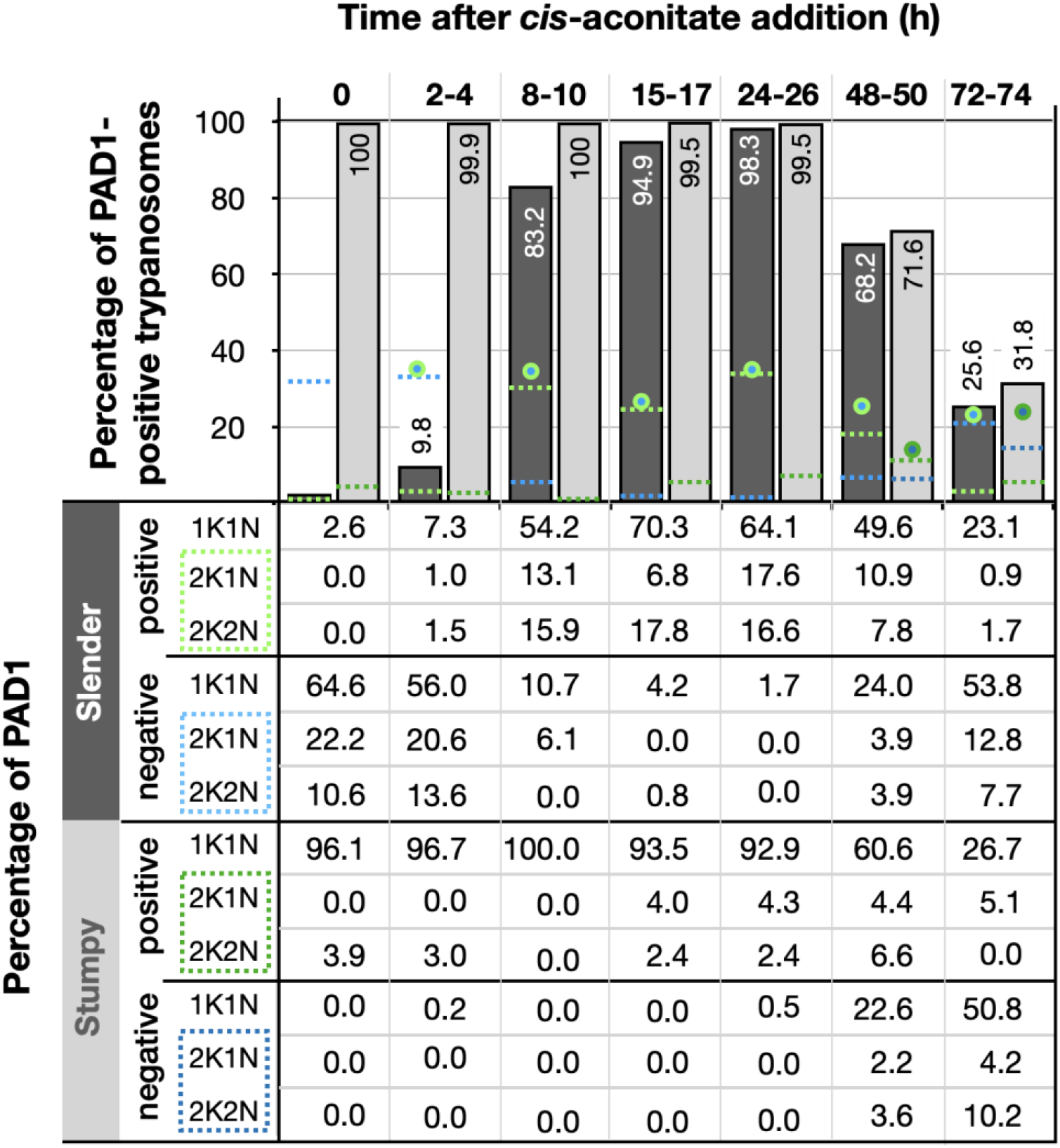
Slender trypanosomes activate the PAD1 pathway *in vitro* without cell cycle arrest. Cultured slender or stumpy trypanosomes were differentiated *in vitro* by the addition of *cis-aconitate* and temperature reduction to 27°C. At the times indicated, trypanosomes were analysed for the expression of the fluorescent stumpy reporter GFP:PAD1^UTR^, as in Fig. 4. Slender cells (n=1653) are shown by dark gray bars and stumpy cells (n=1798) in dark gray bars. Slender and stumpy trypanosomes were also stained with DAPI and the configuration of the nucleus (N) and kinetoplast (K) was microscopically determined to identify the cell cycle stage. Percent of the population as either PAD1-positive and PAD-1 negative in the different cell cycle stages are indicated in the bottom table. The cell cycle stages are also displayed visually in top bar graph. As seen, while the total percentage of dividing slender cells remains constant over time (blue/green circles), the percentage of PAD-1 positive slender cells steadily increases (dotted green line) and the percentage of PAD-1 negative cells steadily decreases (dotted blue line). This shows that slender cells can seamlessly turn on the PAD-1 pathway, without arresting in the cell cycle. Though a small portion of the stumpy population is seen dividing throughout the time points (dark green dotted lines - PAD-1 positive), cells do not return to a normal cell cycle profile until 48 hours after the addition of *cis*-aconitate (total percentage of dividing cells shown as dark blue/green circles). As the stumpy cells become more procyclic, they begin to lose their PAD-1 positive signal and increase in PAD-1 negative dividing cells (dark blue dotted lines). Data were compiled from five independent experiments, with each time point being analysed in at least two separate experiments.

**Figure 7.**
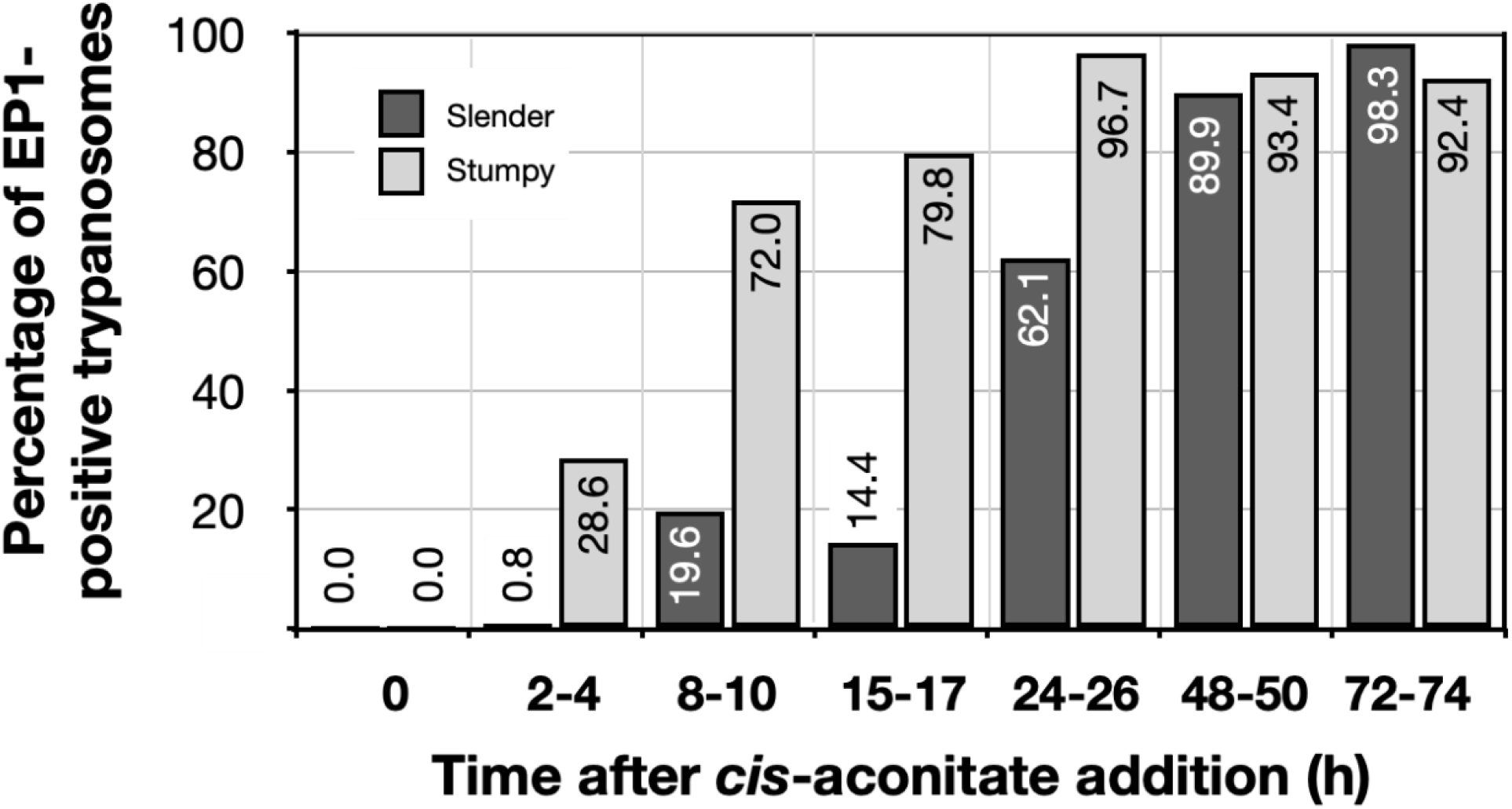
Slender trypanosomes differentiate to the procyclic life cycle stage in vitro without cell cycle arrest. Cultured slender or stumpy trypanosomes were differentiated *in vitro* by the addition of *cis*-aconitate and temperature reduction to 27°C. At the times indicated, trypanosomes were analysed for the expression of the procyclic fluorescent reporters EP1:YFP, as in Fig. 5 Stumpy cells (n=1798) are shown by light gray bars and slender cells (n=1653) by dark gray bars.

By comparison, stumpy parasites (Fig. 6, Stumpy rows) responded to *in vitro cis*-aconitate treatment with rapid expression of the EP1:YFP marker, with 28.6% of all cells being positive within 2-4 hours (Fig. 7). After one day, EP1 was present on almost all (96.7%) stumpy trypanosomes. The cell cycle analysis revealed that the parasites were not dividing, however (Fig. 6, Stumpy rows). The first cells re-entered the cell cycle only after 15-17 hours, and a normal procyclic cell cycle profile was not reached until day 3. Thus, the *in vitro* differentiation supported the *in vivo* observations, demonstrating that pleomorphic slender trypanosomes are able to directly differentiate to the procyclic stage without becoming cell cycle-arrested stumpy cells. The surface expression of EP1 is also of note, as it has been shown that in slender bloodstream parasites, ectopically expressed EP1 does not enter the cell surface, but is retained in endosomes and the flagellar pocket (Engstler & Boshart, 2004). Hence, as in stumpy trypanosomes, the slender trypanosome cell surface access block is lifted by triggering the PAD1 pathway. Furthermore, the overall developmental capacity and differentiation kinetics of both life cycle stages are comparable, *in vitro* and *in vivo*.

## Discussion

Our observations suggest a revised view of the life cycle of African trypanosomes (Fig. 8). We show that one trypanosome suffices to produce robust infections of the tsetse vector, and that the stumpy stage is not essential for tsetse transmission. Slender parasites can complete the complex life cycle in the fly with comparable overall success rates and kinetics as the stumpy stage. Interestingly, the stumpy stage appears more able to establish initial infections in the fly midgut (Table 1, column 5, MG), while slender-derived parasites appear to produce salivary gland infections more efficiently than stumpy-derived counterparts (Table 1, column 8, TI). At first sight, this discrepancy may be related to a greater resistance of the stumpy stage to the digestive environment in the fly’s gut, as has been suggested (Matetovici, De Vooght, & Van Den Abbeele, 2019; Nolan et al., 2000). This, however, is not supported by our data. We have not observed cell death of monomorphic or pleomorphic slender cells in infected tsetse midguts. And even if so, why then should slender-derived cells perform better in the second part of the life cycle? As there will not be a difference between slender- and stumpy-derived procyclic cells, the difference observed must be based on the behavior of bloodstream parasites in the midgut. It is tempting to speculate that one decisive factor could be trypanosome motility. Slender trypanosomes exhibit significantly higher motility compared to stumpy trypanosomes (Bargul et al., 2016). Thus, the mean square displacement in the midgut will be much larger for slender parasites. While stumpy trypanosomes probably never reach the “midgut exit” before differentiation to the insect stage, slender trypanosomes could already be located close to the proventriculus before starting to differentiate to the procyclic stage. Thus, passage through the proventriculus could occur immediately, and the slender-derived trypanosomes could rapidly progress to the mesocyclic stage. This faster mesocyclic progression would result in a less-pronounced infection of the midgut, and a higher TI-value for the slender-derived trypanosomes. While the above hypothesis is consistent with our data, experimental proof would be extremely challenging to obtain. The recent demonstration that glucose levels are a developmental trigger in addition to the well-characterised ones of cold shock and cis-aconitate adds another layer of complexity to the early events during tsetse infection (Qiu et al., 2018).

**Figure 8.**
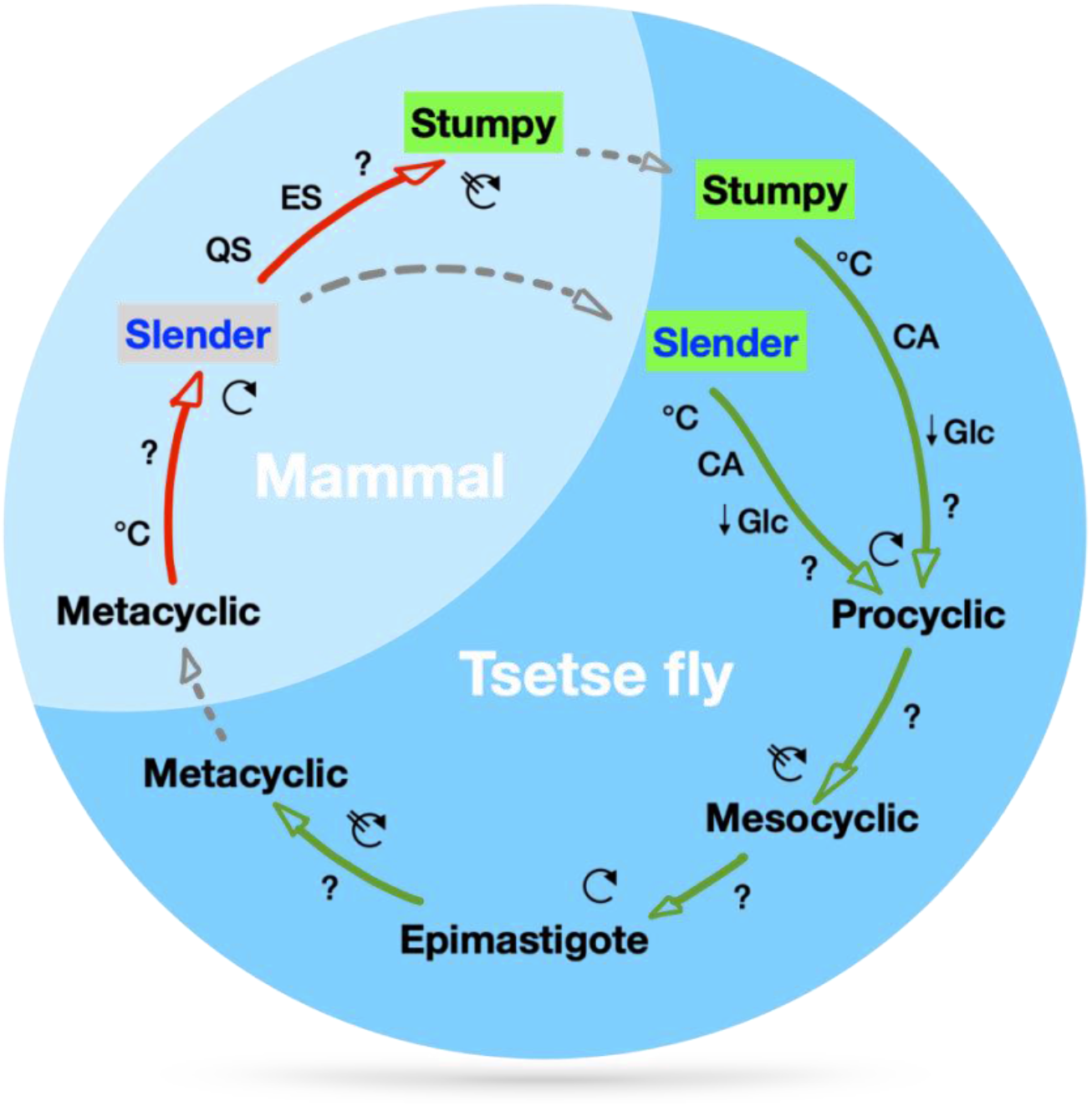
A revised life cycle for the parasite *Trypanosoma brucei.* Cell-cycle-arrested metacyclic trypanosomes are injected by the tsetse fly into the mammalian host’s skin. There, the parasites re-enter the cell cycle, and proliferate as slender forms in the blood, while disseminating into the interstitium and various tissues, including fat, and brain. At least two triggers (SIF or ES) launch the PAD1-dependent differentiation pathway (light green boxes) to the cell cycle-arrested stumpy bloodstream stage. Stumpy trypanosomes can establish a fly infection when taken up with the bloodmeal of a tsetse. This work reveals that proliferating slender stage trypanosomes are equally effective for tsetse transmission, that a single parasite suffices, and that no cell cycle arrest is required for differentiation to the procyclic insect stage.

The dogma that cell cycle-arrested stumpy cells are the only trypanosomes that infect the tsetse fly has never been experimentally challenged, although there are quite a number of reports that point against an exclusive role for stumpy parasites in the life cycle. Koch’s detailed report on the activities of the German sleeping sickness commission sent to East Africa in 1906/7 states that the trypanosome numbers in the blood of human sleeping sickness infections was always very low (Koch, 1909). From his data we have calculated an average blood parasitaemia between 10 and 100 trypanosomes cells/ml (see Supplemental text doc). This means that 2 or fewer trypanosomes would be present in an average tsetse bloodmeal, again highlighting the rarity of a tsetse taking up a stumpy cell. In 1930, Duke discussed the evidence for the essential status of the stumpy stage for tsetse transmission, and his data did not support it (Duke, 1930). Further, Baker and Robertson in 1957 compared the infection capability of *T. rhodesiense* and *T. brucei* using guinea pig feeding (Baker & Robertson, 1957). They concluded: ‘Neither the morphology nor the intensity of the parasitaemia in the infecting mammal was obviously related to the subsequent infection rates in the tsetse-flies.’ In 1990, Bass and Wang, in fact, suggested that the stumpy stage may be dispensable for development to the insect stage (Bass & Wang, 1991). The experiments, however, were in part inconclusive, mainly because a molecular marker for the stumpy stage was missing. The discovery of SIF in the 1990s and the realisation that quorum sensing underpinned the differentiation to the stumpy stage led to an assumption that the slender stage had no role to play in the transmission event. Subsequent research has been focused on the details of stumpy formation, while the developmental role of the stumpy cell has not undergone further examination. The above publications all relate to what is nowadays referred to as the transmission paradox, the persistence and circulation of trypanosomiasis in a population even when parasitaemia levels in individuals are low or close to elimination (Capewell et al., 2019). When parasitaemia is low, stumpy trypanosomes are characteristically absent, making the probability of being ingested by a tsetse fly (which on average ingests 20μl of blood) extremely low. Yet trypanosomiasis persists, even when statistically it should by now have been eliminated. Solutions to the paradox have long been hypothesized and variously include flawed diagnostic testing, asymptomatic cases, and animal reservoirs (Alvar et al., 2020). Recent work using theoretical modelling suggests that for *T. gambiense*, trypanosomes residing in the skin of humans could solve the problem (Capewell et al., 2019). However, there are currently no data available on the number of trypanosomes located in asymptomatic human skin, nor have the kinetics of fly uptake of skin-localised trypanosomes been explored. Also, different tsetse-transmitted trypanosome species reveal rather distinct distributions in the host, such as *Trypanosoma congolense* preferentially residing in small blood vessels (BANKS, 1978). Thus, while trypanosomes in the skin may be important for the persistence of the parasites, their existence alone does not automatically solve the transmission paradox. As tsetse are blood pool feeders, it actually does not matter if the trypanosomes reside in the skin, fat tissue, or blood. Just one or two parasites, stumpy or slender, suffice for infection. Furthermore, stumpy cells will inevitably run into an age-related problem. They are not replicative, and their lifetime is limited to roughly 3 days (Turner et al., 1995). In the fly, re-entry into the cell cycle is by no means immediate, but takes at least one day. Following induction of cell cycle arrest, the stumpy cells would need to be taken up by the fly within one day. Thus, only a subset of rather young stumpy cells would prove successful in the midgut. It is important to note that this is not the case in our experiments, as only freshly differentiated stumpy cells were used for tsetse infection. Thus, our experiments in fact overestimate the success of stumpy stage trypanosomes.

It is worth emphasizing that our data provide a possible solution to the transmission paradox without falsifying any of the extensive published work on stumpy trypanosomes. We have shown that slender and stumpy trypanosomes are equally competent for fly passage. The PAD1 pathway has an essential role in preparing both bloodstream stages for differentiation to the procyclic cell stage. For successful passage through the tsetse fly, however, the stumpy stage is not uniquely required. Along similar lines, it is worth noting that *Trypanosoma congolense*, the principal causative agent of the cattle plague nagana, infects tsetse flies without manifesting a cell cycle-arrested stumpy stage(Rotureau & Van Den Abbeele, 2013). Thus, the essential biological function of the stumpy life cycle stage in *T. brucei* may not be transmission, but rather quorum sensing (SIF)-dependent control of population size in the host. This pathway can be triggered in other ways, and even at low levels of parasitaemia, for example by VSG expression site attenuation (ES)(Zimmermann et al., 2017). The capacity for inducing cell cycle arrest at the single cell level might actually have been important for the evolution of antigenic variation. As not all trypanosome species develop a stumpy life cycle stage (Rotureau & Van Den Abbeele, 2013), density-dependent differentiation at the population level may well be a later innovation in evolution, and specific to the *T. brucei* group. In conclusion, our work exemplifies a high degree of plasticity in the life cycle of an important parasite. It shows that the trypanosome life cycle is not rigid but proposes a revised and less rigid view of the trypanosome life cycle and helps solve a longstanding question in parasitology.

## Methods

### Trypanosome culture

Pleomorphic *Trypanosoma brucei brucei* strain EATRO 1125 (serodome AnTat1.1)(Le Ray, Barry, Easton, & Vickerman, 1977) bloodstream stages were grown in HMI-9 medium (Hirumi & Hirumi, 1989), supplemented with 10% (v/v) fetal bovine serum and 1.1%(w/v) methylcellulose (Sigma 94378, Munich, Germany)(Vassella et al., 2001) at 37°C and 5% CO_2_. Slender stage parasites were maintained at a maximum cell density of 5×10^5^ cells/ml. For cell density-triggered differentiation to the stumpy stage, cultures seeded at 5×10^5^ cells/ml were cultivated for 48 hours without dilution. Pleomorphic parasites were harvested from the viscous medium by 1:4 dilution with trypanosome dilution buffer (TDB; 5 mM KCl, 80 mM NaCl, 1 mM MgSO_4_, 20 mM Na_2_HPO_4_, 2 mM NaH_2_PO_4_, 20 mM glucose, pH 7.6), followed by filtration (MN 615 ¼, Macherey-Nagel, Dueren, Germany) and centrifugation (1,400×*g*, 10 min, 37°C)(Zimmermann et al., 2017). Monomorphic *T. brucei* 427 MITat 1.2 13-90 bloodstream stage (Wirtz, Leal, Ochatt, & Cross, 1999) were grown in HMI-9 medium (Hirumi & Hirumi, 1989), supplemented with 10% (v/v) fetal bovine serum at 37°C and 5% CO_2_.

For *in vitro* differentiation to the procyclic insect stage, bloodstream stage trypanosomes were pooled to a cell density of 2×10^6^ cells/ml in DTM medium with 15% fetal bovine serum immediately before use (Overath, Czichos, & Haas, 1986). *Cis*-aconitate was added to a final concentration of 6 mM (Brun et al., 1981; Overath et al., 1986) and temperature was adjusted to 27°C. Procyclic parasites were grown in SDM79 medium (Brun & Schönenberger, 1979), supplemented with 10% (v/v) fetal bovine serum (Hirumi & Hirumi, 1989) and 20 mM glycerol (Schuster et al., 2017; Vassella et al., 2000).

### Genetic manipulation of trypanosomes

Transfection of pleomorphic trypanosomes was done as previously described (Zimmermann et al., 2017), using an AMAXA Nucleofector II (Lonza, Basel, Switzerland). Transgenic trypanosome clones were selected by limiting dilution in the presence of the appropriate antibiotic. The GFP:PAD1^UTR^ reporter construct (Zimmermann et al., 2017) was used to transfect AnTat1.1 trypanosomes to yield the cell line ‘SIF’. The trypanosome ‘ES’ line was described previously (Zimmermann et al., 2017). It contains the reporter GFP:PAD1^UTR^ construct and an ectopic copy of VSG gene MITat 1.6 under the control of a tetracycline-inducible T7-expression system. The EP1:YFP construct was integrated into the EP1-procyclin locus as described previously (Engstler & Boshart, 2004).

### Immunofluorescence

Cells were harvested as stated above, concentration was measured using a Neubauer chamber, and 10^6^ cells per coverslip were taken. The cells were transferred to a 1.5ml tube, washed twice with 1ml of phosphate buffered saline (PBS), resuspended in 500ul of PBS, and fixed by addition of formaldehyde to a final concentration of 4% at room temperature (RT) for 20 minutes (min). The cells were pelleted by centrifugation (750 *xg*, RT, 10min), supernatant removed, resuspended in PBS, and transferred to poly-L-lysine coated coverslips in a 24-well plate. Cells were attached to coverslips by centrifugation (750 *xg*, RT, 4 min). Cells were either permeabilized with 0.25% TritonX-100 in PBS (RT, 5min) and subsequently washed twice with PBS or not permeabilized, so as to allow only surface labelling. Cells were then blocked with 3% BSA in PBS (RT, 30 min), followed by incubation with the primary (1:100 rabbit anti-PAD1; 1:500 IgG1 mouse anti-Trypanosoma brucei procyclin, Ascites, Clone TBRP1/247, CEDARLANE, Ontario, Canada) and secondary antibodies (Alexa488- and Alexa 594-conjugated anti-rabbit and anti-mouse, 1:100, ThermoFisher Scientific, Massachusetts, USA) diluted in PBS (1h, RT for each), with three PBS wash steps after each incubation. After the final wash, coverslips were rinsed with ddH2O, excess fluid removed by wicking, and mounted on glass slides using antifade mounting media with DAPI (Vectashield, California, USA).

### Tsetse maintenance

The tsetse fly colony (*Glossina morsitans morsitans*) was maintained at 27°C and 70% humidity. Flies were kept in Roubaud cages and fed 3 times a week through a silicone membrane, with prewarmed, defibrinated, sterile sheep blood (Acila, Moerfelden, Germany).

### Fly infection and dissection

Teneral flies were infected 1-3 days post-eclosion during their first meal. It is known that teneral flies (flies that are newly hatched and unfed) are more susceptible to midgut infections compared to older flies, and it is an accepted practice in the field to use teneral flies for infections. While all of our infections were done during the flies’ first bloodmeal, it is of note that 1-3 days is rather old for teneral flies (Walshe, Lehane, & Haines, 2011; Wijers, 1958). Depending on the experiment, trypanosomes were diluted in either pre-warmed TDB or sheep blood. For infections with low parasite number (Table 1), the cell density of either stumpy or slender trypanosomes was calculated and the dilutions made directly in blood without harvesting the cells. The infective meals were supplemented with 60 mM N-acetylglucosamine (Peacock, Ferris, Bailey, & Gibson, 2006). For infection with 2400 monomorphic parasites per bloodmeal, cells were additionally treated for 48 hours with 12.5 mM glutathione (GSH) (MacLeod, Maudlin, Darby, & Welburn, 2007) and 100 μM 8-pCPT-cAMP (cAMP) (Vassella et al., 1997).

Tsetse infection status was analyzed between 35 and 40 days post-infection. Flies were euthanized with chloroform and dissected in PBS. Intact tsetse alimentary tracts were explanted and analysed microscopically, as described previously (Schuster et al., 2017). For the analysis of early trypanosome differentiation *in vivo*, slender or stumpy trypanosomes at a concentration of 6×10^5^ cells/ml were resuspended in TDB to the required final concentration and fed to flies. The numbers of flies used and the number of independent experiments carried out are indicated in the figure legends. Results are presented as sample means.

### Fluorescence microscopy and video acquisition

Live trypanosome imaging was performed with a fully automated DMI6000B widefield fluorescence microscope (Leica microsystems, Mannheim, Germany), equipped with a DFC365FX camera (pixel size 6.45 μm) and a 100x oil objective (NA 1.4). For high-speed imaging, the microscope was additionally equipped with a pco.edge sCMOS camera (PCO, Kelheim, Germany; pixel size 6.5 μm). Fluorescence video acquisition was performed at frame rates of 250 fps. For visualization of parasite cell cycle and morphology, slender and stumpy trypanosomes were harvested and incubated with 1 mM AMCA-sulfo-NHS (Thermo Fisher Scientific, Erlangen, Germany) for 10 minutes on ice. Cells were chemically fixed in 4% (w/v) formaldehyde and 0.05% (v/v) glutaraldehyde overnight at 4°C. DNA was visualised with 1 μg/ml DAPI immediately before analysis.

3D-Imaging was done with a fully automated iMIC widefield fluorescence microscope (FEI-TILL Photonics, Munich, Germany), equipped with a Sensicam qe CCD camera (PCO, Kelheim, Germany; pixel size 6.45 μm) and a 100x oil objective (NA 1.4). Deconvolution of image stacks was performed with the Huygens Essential software (Scientific Volume Imaging B.V., Hilversum, Netherlands). Fluorescence images are shown as maximum intensity projections of 3D-stacks in false colours with green fluorescence in green and blue fluorescence in grey.

### Scanning electron microscopy

Explanted tsetse alimentary tracts were fixed in Karnovsky solution (2% formaldehyde, 2.5% glutaraldehyde in 0.1M cacodylate buffer, pH 7.4) and incubated overnight at 4°C. Samples were washed 3 times for 5 minutes at 4°C with 0.1M cacodylate buffer, pH 7.4, followed by incubation for 1 hour at 4°C in post-fixation solution (2.5% glutaraldehyde in 0.1M cacodylate buffer, pH 7.4). After additional washing, the samples were incubated for 1 hour at 4°C in 2% tannic acid in cacodylate buffer, pH 7.4, 4.2% sucrose, and washed again in water (3x for 5 minutes, 4°C). Finally, serial dehydration in acetone was performed, followed by critical point drying and platinum coating. Scanning electron microscopy was done using the JEOL JSM-7500F field emission scanning electron microscope (JEOL, Freising, Germany).

## Data Availability

All datasets generated during this project are provided as online source data. The cell lines used are available upon request.

## Acknowledgements

We thank Nicola Jones, Susanne Kramer, Manfred Alsheimer, Christian Janzen and Ricardo Benavente for discussion and critical reading of the manuscript. We thank Keith Matthews (Edinburgh) for the anti-PAD1 antibody. BM is supported by DFG grant number 396187369. ME is supported by DFG grants EN305, SPP1726 (Microswimmers – From Single Particle Motion to Collective Behaviour), GIF grant I-473-416.13/2018 (Effect of extracellular *Trypanosoma brucei* vesicles on collective and social parasite motility and development in the tsetse fly) and GRK2157 (3D Tissue Models to Study Microbial Infections by Obligate Human Pathogens). ME is a member of the Wilhelm Conrad Roentgen Center for Complex Material Systems (RCCM).

## Author contributions

S.S. designed the experiments, performed the experiments, analysed the data, interpreted the results and wrote the manuscript. I.S. designed the experiments, performed the experiments, analysed the data and interpreted the results. J.L. designed the experiments, performed the experiments, analysed the data, interpreted the results and wrote the manuscript. H.Z., designed the experiments, performed the experiments, analysed the data and interpreted the results. C.R. designed the experiments, performed the experiments, analysed the data and interpreted the results. B.M. interpreted the results and wrote the manuscript. M.E. conceived the study, designed the experiments, analysed the data, interpreted the results and wrote the manuscript.

## Supplementary Video 1

**Supplemental video 1.**
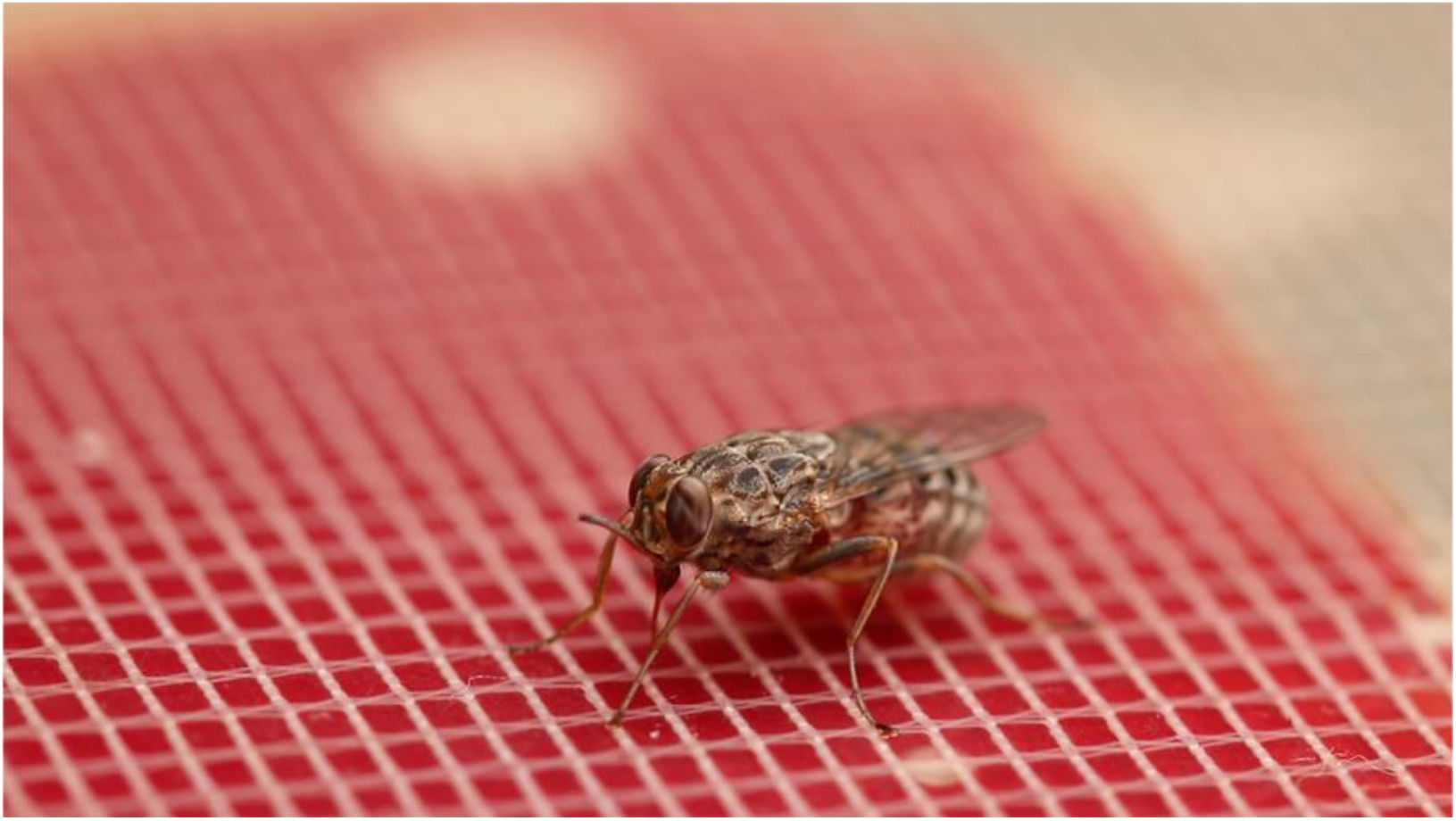
Video of a tsetse fly taking a bloodmeal through

## Supplementary Video 2

**Supplemental video 2.**
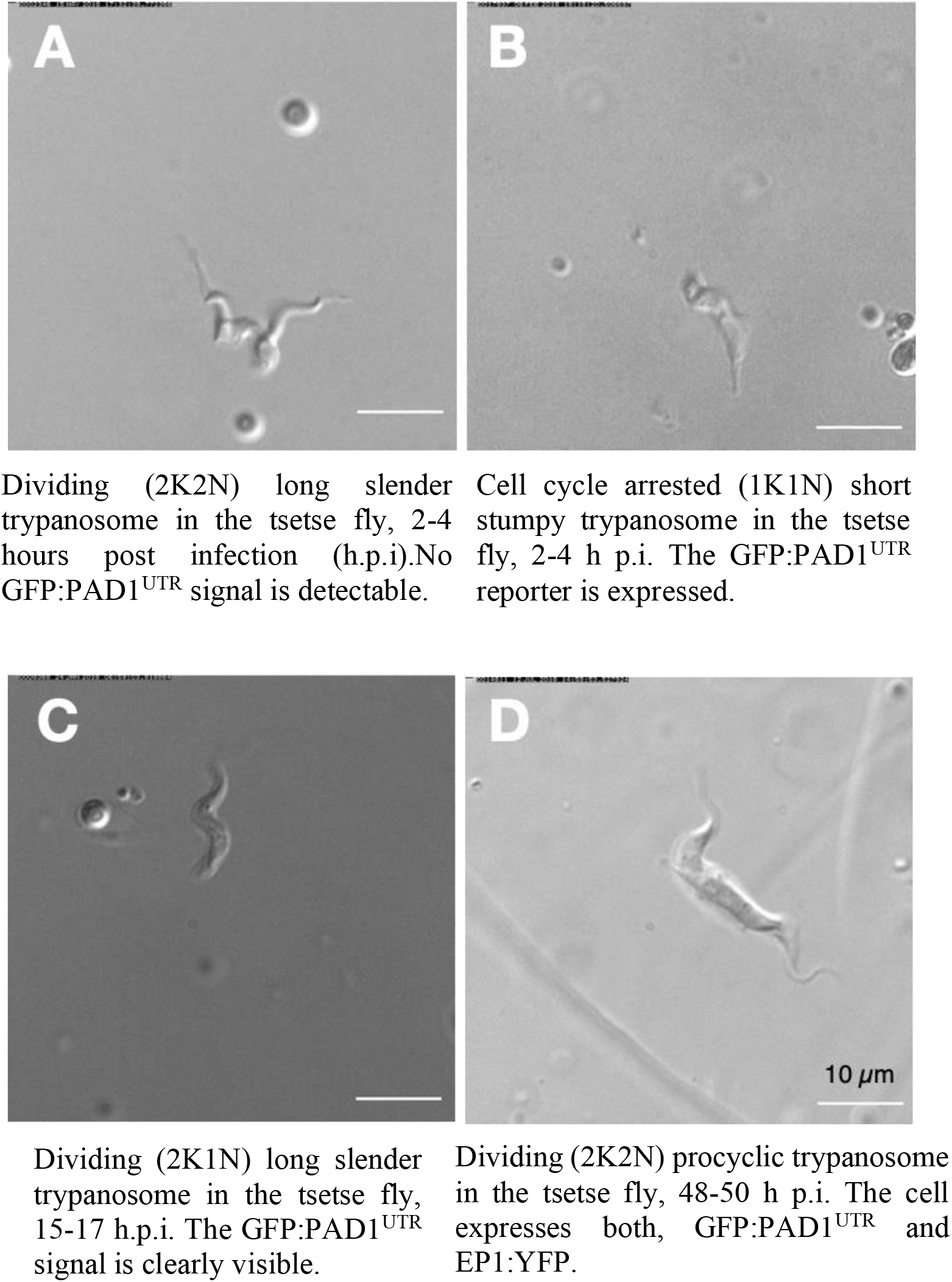
After uptake by the tsetse fly, slender trypanosomes promptly activate the PAD1 pathway, without arresting in the cell cycle. All videos were recorded at 250 fps, and the cell cycle position is indicated by DAPI staining.

## Supplemental Figure 1

**Supplemental figure 1.**
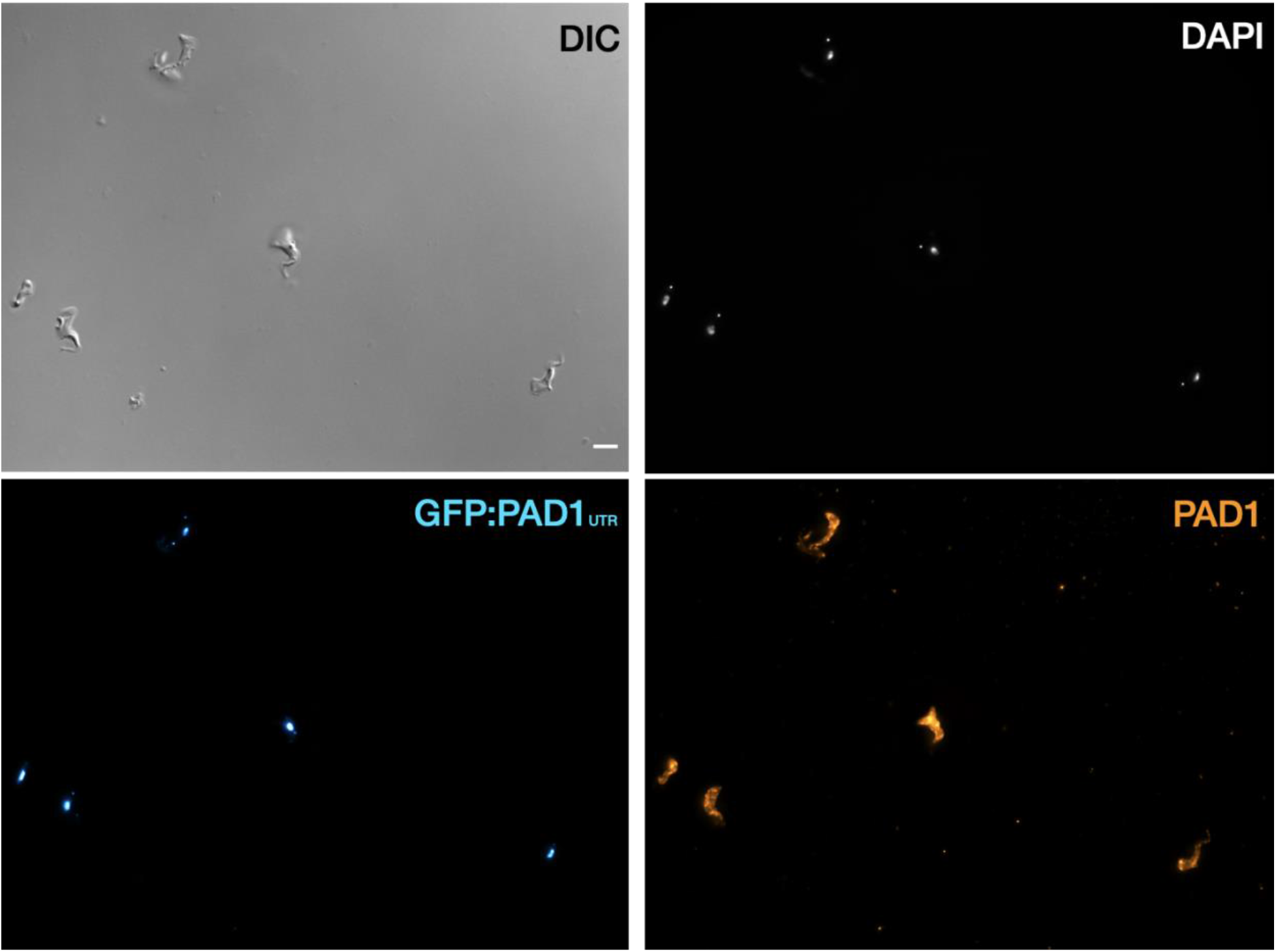
Stumpy trypanosomes express PAD1 on their surface when the GFP:PAD1^UTR^ is expressed. Immunofluorescence using an anti-PAD1 antibody with stumpy trypanosomes (generated with SIF) from the GFP:PAD1^UTR^ cell line. Cells were fixed in formaldehyde and labelled with an anti-PAD1 antibody, without membrane permeabilization, in order to only detect surface-localized proteins. DAPI (gray), GFP:PAD1^UTR^ signal (cyan), and PAD1 protein (orange) are shown. Scale bar = 5 μm.

## Supplemental Figure 2

**Supplemental figure 2.**
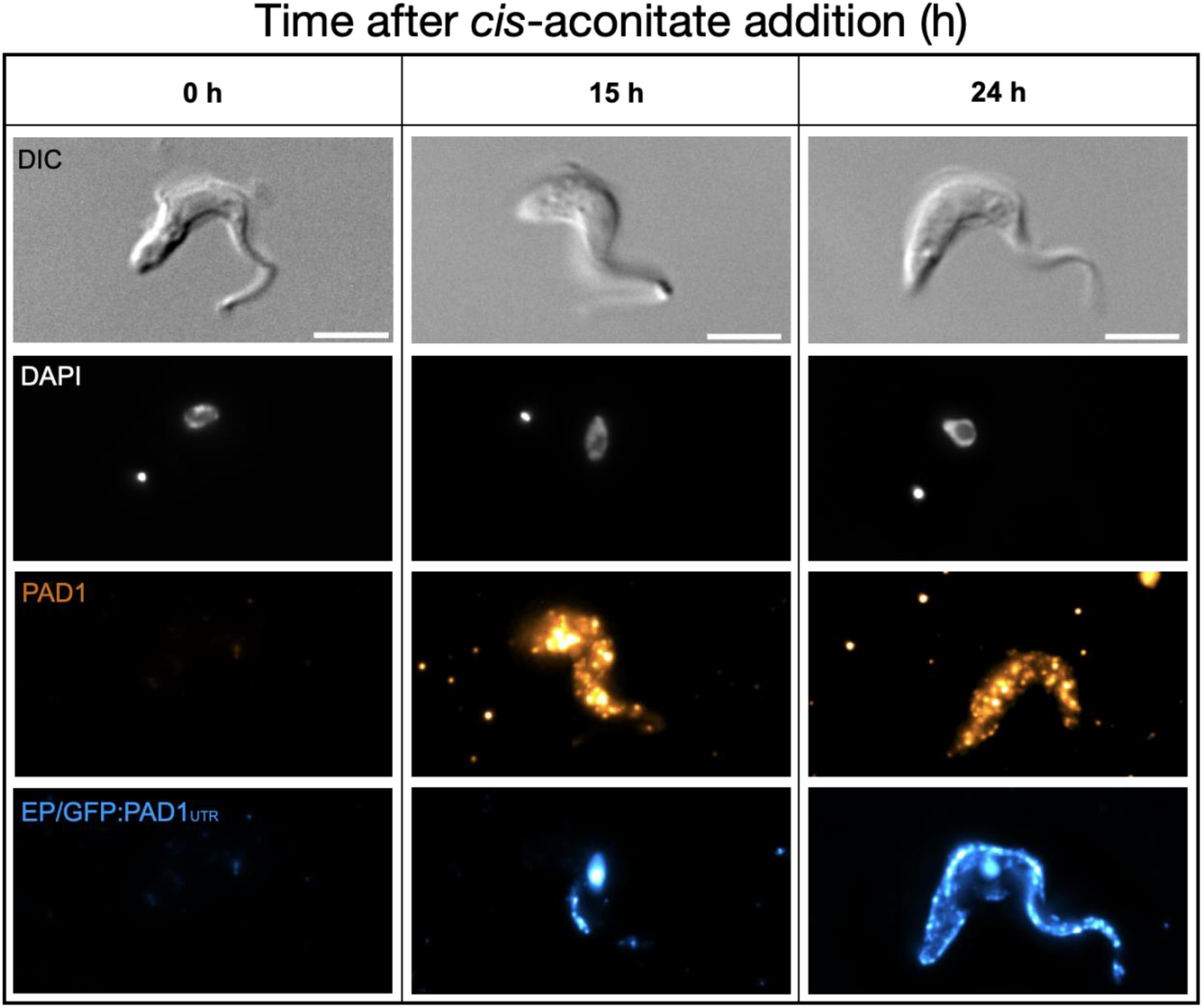
Slender cells express PAD and EP on their surface after the addition of cis-aconitate. Immunofluorescence time course (h = hours) of slender cells from the cell line GFP:PAD1^UTR^ after the addition of cis-aconitate. Cells were fixed in formaldehyde and labelled with both anti-PAD1 and anti-EP antibodies, without membrane permeabilization, in order to only detect surface-localized proteins. The fluorescent nucleus seen in the 15 and 24 hr timepoints are from the background GFP:PAD1^UTR^ signal (containing a nuclear localization sequence and having the same fluorescent signal as the secondary antibody used to target anti-EP), further showing that the PAD-1 mRNA signal is here representative of PAD1 protein on the surface. Dapi (gray), PAD-1 (orange) and EP (blue) are shown. Scale bar = 5 μm.

## Supplemental Text 1.

**Supplementary information on the number of trypanosomes in human blood samples, based on original observations reported by Robert Koch in 1906/07**

**(Translated with DeepL Pro)**

**Robert Koch. Report on the activities of the commission sent to East Africa to research sleeping sickness in 1906/07 (Verlag von Julius Springer, Berlin, 1909)**

(page 17) “If now sick people, in whose blood trypanosomes can be detected, are examined quite carefully and daily, as we have done several times, then one first learns that the number of trypanosomes in the blood is almost always very low. Often only one or two trypanosomes are present in a preparation containing several drops of blood. Five to ten trypanosomes in one preparation are already a rather rich yield. We have only exceptionally seen a larger number of trypanosomes, so that every second to third field of view of the very thick layer of the preparation was filled with one trypanosome. Such quantities of trypanosomes, which are almost regularly seen in the blood of laboratory animals, have never been found in the blood of humans.

The occurrence of trypanosomes in the blood is quite irregular. If they were found for one or a few days, they suddenly disappear and usually stay away for 2 to 3 weeks, only to reappear again.

They are then very sporadic at the beginning, become a little more numerous the next day and maybe even the third day, then decrease again for one or two days and disappear again. It seems as if they appear periodically in the blood, their presence lasts 2 to 5 days and their absence 2 to 3 weeks. In most cases, the recurrence of trypanosomes is associated with an increase in temperature and increased symptoms of disease, especially headaches and chest pain.

It is necessary to be familiar with the periodic appearance of trypanosomes in the blood in order not to make too many futile examinations during the diagnostic examination of the blood.

In blood preparations, trypanosomes have a very different appearance depending on whether they lie on the band or more towards the inside. On the periphery they appear in terms of their size, the shape of the nucleus, visibility of the undulating membrane and the flagella, just as one is used to see them in smear preparations of the blood of the test animals. But in the thick layers of the inner parts of the preparation they look considerably smaller, their colour is darker, they also have a rounded appearance, the nucleus is smaller, membrane and flagellum are hardly visible, often they seem to be missing. However, this different appearance is not due to the different composition of the trypanosomes, but is only caused by the preparation. At the edges they dry up in a very thin layer and very fast. They are thus spread out, stretched to a certain extent and immediately fixed in this form by drying. In the thick blood layer of the preparation, the drying process is only gradual, leaving the Trypanosoma time to dry in its original cylindrical shape with more or less strong shrinking of the whole body and especially of the undulating membrane and the flagella.”

**(Original Text in German)**

**Robert Koch. Bericht über die Tätigkeit der zur Erforschung der Schlafkrankheit im Jahre 1906/07 nach Ostafrika entsandten Kommission (Verlag von Julius Springer, Berlin, 1909)**

(Seite 17) „Wenn nun Kranke, in deren Blut Trypanosomen nachzuweisen sind, recht sorgfaltig und täglich untersucht werden, wie wir das des öfteren getan haben, dann erfährt man zunächst, daß die Anzahl der Trypanosomen im Blute fast immer eine sehr geringe ist. Auf ein Präparat, welches mehrere Tropfen Blut enthält, kommen oft nur ein oder zwei Trypanosomen. Fünf bis zehn Trypanosomen in einem Präparat bilden schon eine ziemlich reiche Ausbeute. Wir haben nur ausnahmsweise eine größere Zahl von Trypanosomen gesehen, so daß auf jedes zweite bis dritte Gesichtsfeld der sehr dicken Präparatenschicht ein Trypanosoma kam. Solche Mengen von Trypanosomen, wie man sie fast regelmäßig im Blute der Versuchstiere zu sehen bekommt, haben wir niemals im Blute der Menschen angetroffen.

Das Vorkommen der Trypanosomen im Blute ist ziemlich unregelmäßig. Wenn sie einen oder einige Tage lang gefunden wurden, dann sind sie plötzlich verschwunden und bleiben gewöhnlich 2 bis 3 Wochen fort, um dann wieder zum Vorschein zu kommen.

Sie sind dann anfangs ganz vereinzelt, werden am nächsten und vielleicht auch noch am dritten Tage ein wenig zahlreicher, nehmen dann wiederum ein bis zwei Tage ab und verschwinden von neuem. Es hat den Anschein, als ob sie periodenweise im Blute erscheinen, und zwar dauert ihr Vorhandensein 2 bis 5 Tage und ihr Fehlen 2 bis 3 Wochen. Meistens sind mit dem Wiederauftreten der Trypanosomen eine Temperatursteigerung und verstärkte Krankheitssymptome, namentlich Kopf-und Brustschmerzen, verbunden.

Man muß mit dem periodenweisen Erscheinen der Trypanosomen im Blute vertraut sein, um bei der diagnostischen Untersuchung des Blutes nicht zu viele vergebliche Untersuchungen zu machen.

In den Blutpräparaten haben die Trypanosomen ein sehr verschiedenes Aussehen, je nachdem sie am Bande oder mehr nach dem Innern zu liegen. Am Rande erscheinen sie in bezug auf ihre Größe, auf die Gestalt des Kerns, Sichtbarkeit der undulierenden Membran und der Geißel, ebenso wie man sie in Ausstrichpräparaten vom Blut der Versuchstiere zu sehen gewohnt ist. Aber in den dicken Schichten der inneren Partien des Präparates sehen sie erheblich kleiner aus, ihre Farbe ist dunkler, sie haben auch ein rundliches Aussehen, der Kern ist kleiner, Membran und Geißel sind kaum zu er-kennen, oft scheinen sie zu fehlen. Dieses verschiedene Aussehen beruht nun aber nicht auf verschiedener Beschaffenheit der Trypanosomen, sondern ist nur durch die Präparation bedingt. Am Rande trocknen sie in sehr dünner Schicht und sehr schnell ein. Dabei werden sie also der Fläche nach ausgebreitet, gewissermaßen gestreckt und in dieser Form durch das Eintrocknen sofort fixiert. In der dicken Blutschicht des Präparats geht der Eintrocknungsprozeß nur allmählich vor sich, und da bleibt dem Trypanosoma Zeit, in seiner ursprünglichen walzenförmigen Gestalt unter mehr oder weniger starkem Schrumpfen des ganzen Körpers und ganz besonders der undulierenden Membran und der Geißel zu trocknen.”

**Based on Koch’s observations, the following estimations can be made:**

One drop of blood =**50 μl** and “several drops” are 5 drops = 250 μl; maximum count was 5-10 trypanosomes per 5 drops on average, which means 20-40 trypanosomes are present in one milliliter of blood. Hence, one tsetse bloodmeal of 20 μl would contain 0.4 to 0.8 trypanosomes.

One drop of blood =**20 μl** and “several drops” are 5 drops = 100 μl; maximum count was 5-10 trypanosomes per 5 drops on average, which means 50-100 trypanosomes per ml are present in one milliliter of blood. Hence, one tsetse bloodmeal of 20 μl would contain 1 to 2 trypanosomes.

